# From wing shape to fluid dynamics in the barn owl (*Tyto alba*)

**DOI:** 10.1101/2020.05.29.121178

**Authors:** Jorn A Cheney, Jonathan PJ Stevenson, Nick E Durston, Jialei Song, Masateru Maeda, Shane P Windsor, James R Usherwood, Richard J Bomphrey

## Abstract

Birds morph their wings and tail in order to glide under a wide range of aerodynamic conditions. Gross wing morphing has been described in a multitude of studies, but the finer details of wing morphing are still unknown. Here, we measured the changes in wing shape and pose in a barn owl, *Tyto alba*, when gliding across a range of fifteen self-selected speeds. We found that *T. alba* does not use fine-wing shape control to glide at slow speeds in steady conditions, with the measured wing shapes being highly consistent across all flights. Instead, *T. alba* relied upon wing postural control (gross pitch) and changes in both tail shape and pose to modulate aerodynamic force. A consistent wing shape provides an exceptional aerodynamic tool for understanding gliding flight in birds through postural change and tail morphing. This geometry was used as the basis for computational fluid dynamics simulations which gave very similar wake measurements and weight support to those measured in flight. This geometry is provided here to assist other researchers interested in exploring the fluid dynamics behind gliding flight in birds.

## Introduction

Birds fly over a wide range of speeds by morphing their wings and tail to modulate aerodynamic force production. At the fastest gliding speeds, birds hold their wings in a folded and swept back configuration, and the tail is held in a contracted state to minimise area; as in the stooping postures of hunting birds (Ponitz et al., 2014). With decreasing speed, birds unfold their wings, reduce wing sweep until they spread laterally, but maintain the tail in the same contracted state (Rosen and Hedenström, 2001; Henningson and Hedenström, 2011). Finally, at the slowest speeds, the wings approach a fully unfolded state, and the tail spreads to augment lift generation (Pennycuick, 1968; Tucker, 1992; Rosen and Hedenström, 2001; Henningson and Hedenström, 2011).

The wing morphing that occurs in flight is critical for expanding the flight envelope of birds. We separate wing morphing into two mechanisms: changes in posture, and changes in shape. Changes in posture can be described by rigid-body transformations; *i.e*., the wing rotating about the shoulder or the tail pitching up and down. Changes in shape are deformations, such as changes in wing or tail area or camber. Both mechanisms are critical for minimising drag, increasing manoeuvrability, and modifying stability.

The full capacity for wing-shape change in gliding birds is unknown due to difficulty of measurement. Wings dramatically morph during flapping flight (Wolf and Konrath, 2015), but it is unclear how many active degrees of freedom are available for controlling morphing. Bird wings are highly coupled; the internal structure of a bird’s wing acts as a linkage, where the bones and feathers generally move together in prescribed ways (Matloff et al., 2020; Harvey et al., 2019), and much of the obvious shape change is controlled by a single degree of freedom. However, there is a theoretical capacity for near-infinite degrees of freedom in wing shape due to anatomical complexity. Bird wings are multi-jointed, flight feathers are actuated by tonic muscles (Hieronymus, 2016), ligament linkages are elastic, and aeroelasticity passively bends feathers into new shapes. Here, we aim to examine the magnitude and scope of smaller, more subtle, wing shape change across a band of flight speeds.

We examine how a barn owl, *Tyto alba*, changes wing shape and/or pose when gliding across a range of self-selected speeds. We measured wing shape using photogrammetric methods (Fig 1; similar to those in Durston et al., 2019), which provided accurate three-dimensional geometry of the entire bird. The specific aerodynamic function of shape and pose change is beyond the scope of this study; however, we highlight a few important relevant morphological traits related to stability and control. For finer analysis, we provide a geometrical model of the bird for future work to address specific aerodynamic function using computational fluid dynamics. Further, we have validated the computational output from this model against the downwash distribution measured in live birds using particle-tracking velocimetry, and by comparing the force generated to weight support. Results of our shape analysis suggest that this geometrical model is representative of flight in the slow-moderate speed range.

**Figure 1.**
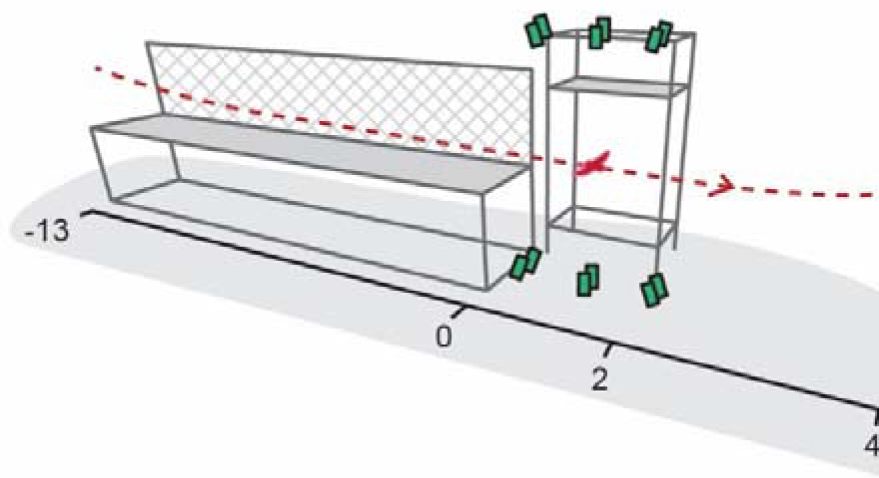
View of indoor flight corridor. The flight path (red dashed line) is constrained by the building wall (not shown) and a mesh wall (hatched). Six cameras (green boxes) record images of the surface of the bird for reconstruction.

## Results

### Camera calibration accuracy

We estimate reconstruction error attributed to camera calibration by comparing a high-accuracy laser-scan of a physical model bird to that of the point cloud reconstruction. Median unsigned error was 0.6 mm; 72% of all points were within 1 mm from the laser-scan, and 95% of all points were within 2.2 mm (Fig 2).

**Figure 2.**
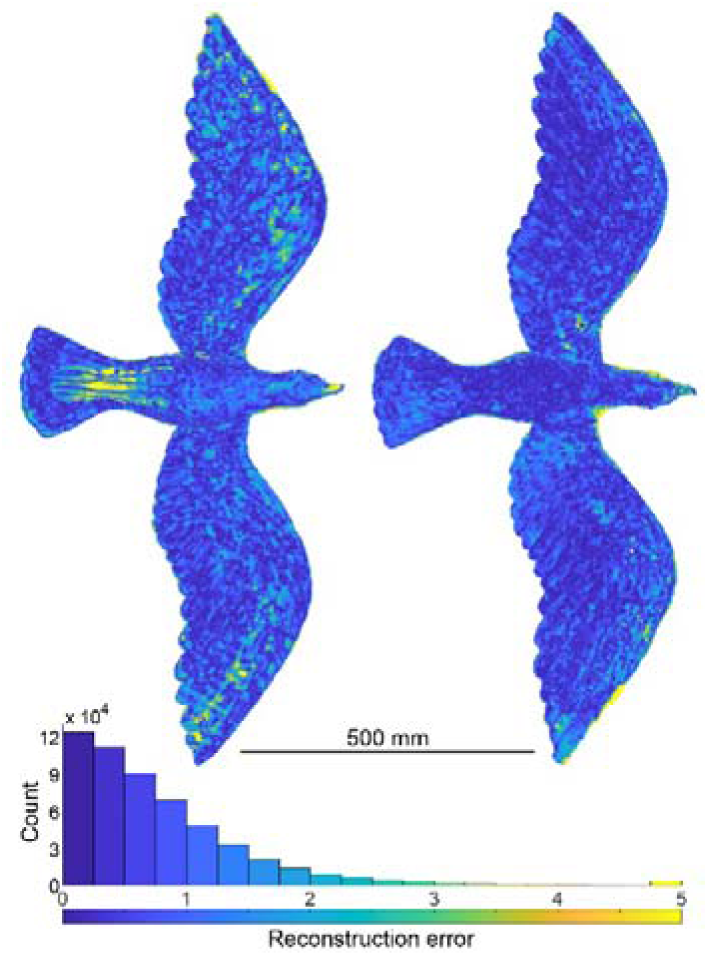
Error in reconstruction of a rigid laser-scanned bird model. Reconstructed point cloud is color-coded based on absolute error. Histogram below shows that the error is generally sub-millimetre. Reconstruction performed with identical camera and light placement used during data collection.

### Self-selected speeds and statistical analysis

Across fifteen glides, flight speed was 7.7 ± 0.4 m·s^-1^ (mean ± standard deviation; n=15). One flight was notably slower at 6.6 m·s^-1^. To account for this in our regression analysis, we used bootstrap statistical methods, and computed regressions against speed with resampled residuals to reduce the likelihood that this single flight disproportionately affected the results.

### Accelerations

Accelerations were generally low (Table 1). Average upward acceleration was 0.08 ± 0.09 g (g = 9.82 m·s^-2^)and forward acceleration was -0.06 ± 0.08 g. There was a significant but weak trend between forward acceleration and speed, with speed accounting for 20% of the variance (r^2^) in forward acceleration (r: mean ± s.e.m., -0.45 ± 0.19). Note, a glide of average speed was an outlier with large upward acceleration, 0.32 g, deviating by 2.6 standard deviations from the mean.

**Table 1.** Distribution of body dynamics of T. alba across the fifteen glides.

| | mean ± standard<br>deviation | observed<br>range |
| --- | --- | --- |
| Speed ( m s<sup>-1</sup> ) | 7.7 ± 0.4 | [6.6, 8.1] |
| Glide angle (deg) | 1.9 ± 1.0 | [0.0, 3.7] |
| Body Pitch (deg) | 2.8 ± 0.9 | [1.1, 4.0] |
| Acc. Forward ( m s<sup>-2</sup> ) | -0.57 ± 0.80 | [-2.0, 0.9] |
| Acc. Vertical ( m s<sup>-2</sup> ) | 0.75 ± 0.92 | [-0.51, 3.10] |

### Insight into speed range: relationship between speed and tail pose and shape

Tail pose and shape varied with speed. The tail was always pitched with the tip of the tail below the root, but with increasing speed, the tail flattened and the angle relative to the torso decreased (mean ± s.e.m., slope: -5.0 ± 0.9 degrees per m·s^-1^; r: -0.71 ± 0.24; Fig 3A). Tail shape changed with increasing speed through area reduction (slope: -17.8 ± 7.0 cm^2^ per m·s^-1^; r: -0.46 ± 0.31; Fig 3B), which can be accounted for by angular reduction in spread of the tail feathers (Fig 3C).

**Figure 3.**
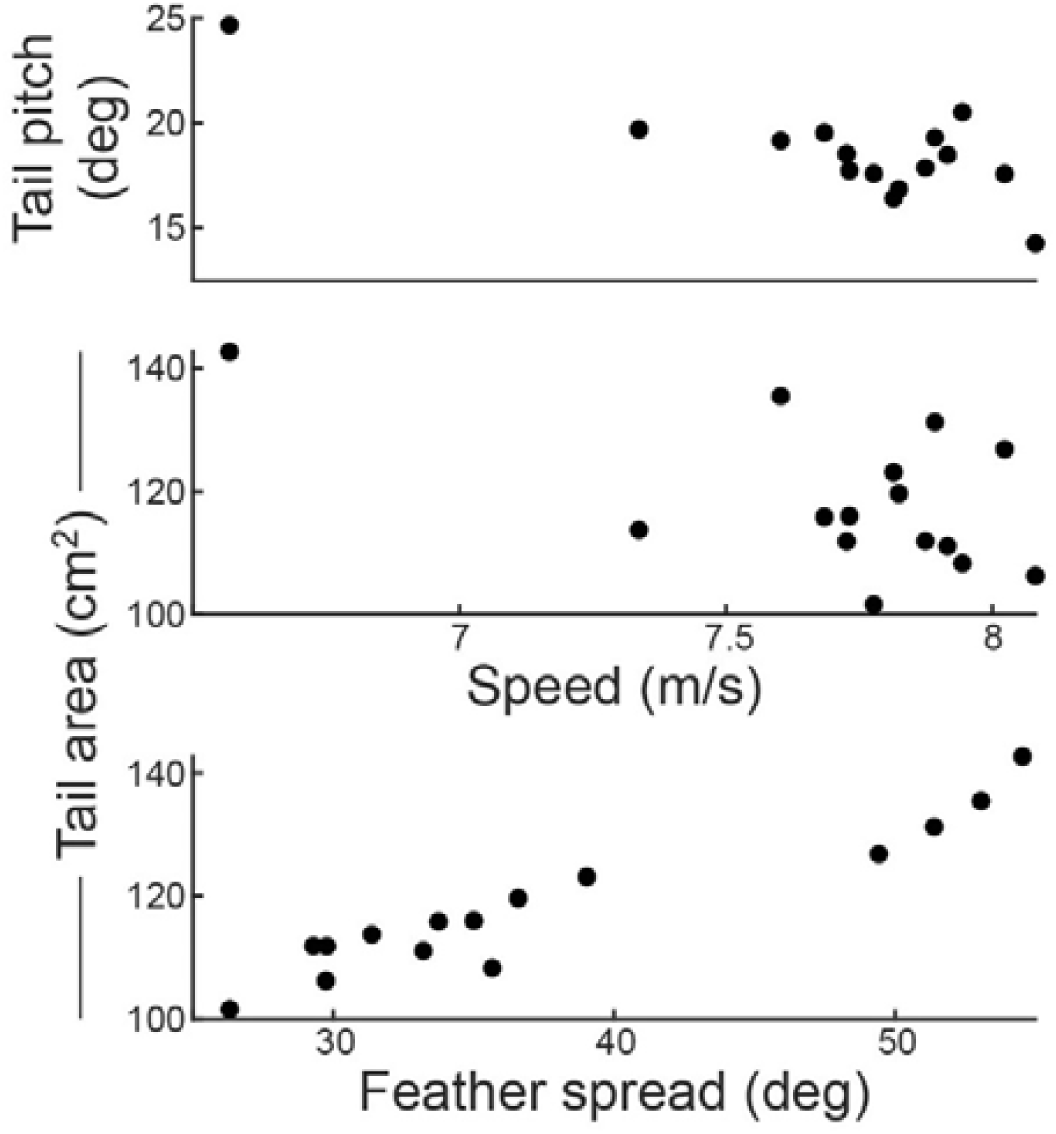
Tail dynamics vary with speed. Tail pitch and area both decreased with increasing flight speed. Tail area changes size in response to feather spread angle.

### Similarity in spanwise distributions of wing shape

After accounting for rigid-body translations and rotations of the left and right wings, wing shape did not vary dramatically across flights. The wings are similar in profile (Fig 4); across planform: camber and thickness (Fig 5), and across span: twist--change in chord angle per span, chord length, and relative position of the chords—as expressed as curvature of the line tracing the quarter-chord (Fig 6). The standard deviation in all metrics is low. Variation does increase slightly at the wingtips for metrics normalised by chord length such as chord-pitch angle, camber, and thickness, but this is likely attributable to chord length approaching negligible length.

**Figure 4.**
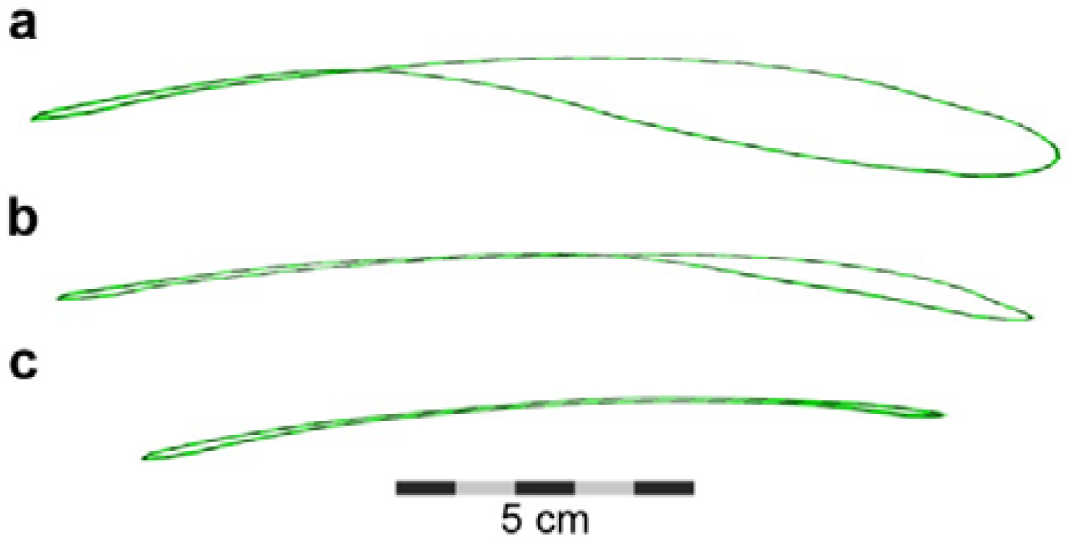
Mean wing profile and standard error at three sections. Wing shape is well described by the mean (black dashed line), as standard error (green shaded area; n= 30) is VERY low. We use a dashed line for the mean profile, to allow the standard error to be visualized at a) 25% wing length; b) 50% wing length; c) 75% wing length. We compute and display standard error independently on both axes. The upper and lower surfaces do cross, suggesting a small amount of systematic error exists in our measurements.

**Figure 5.**
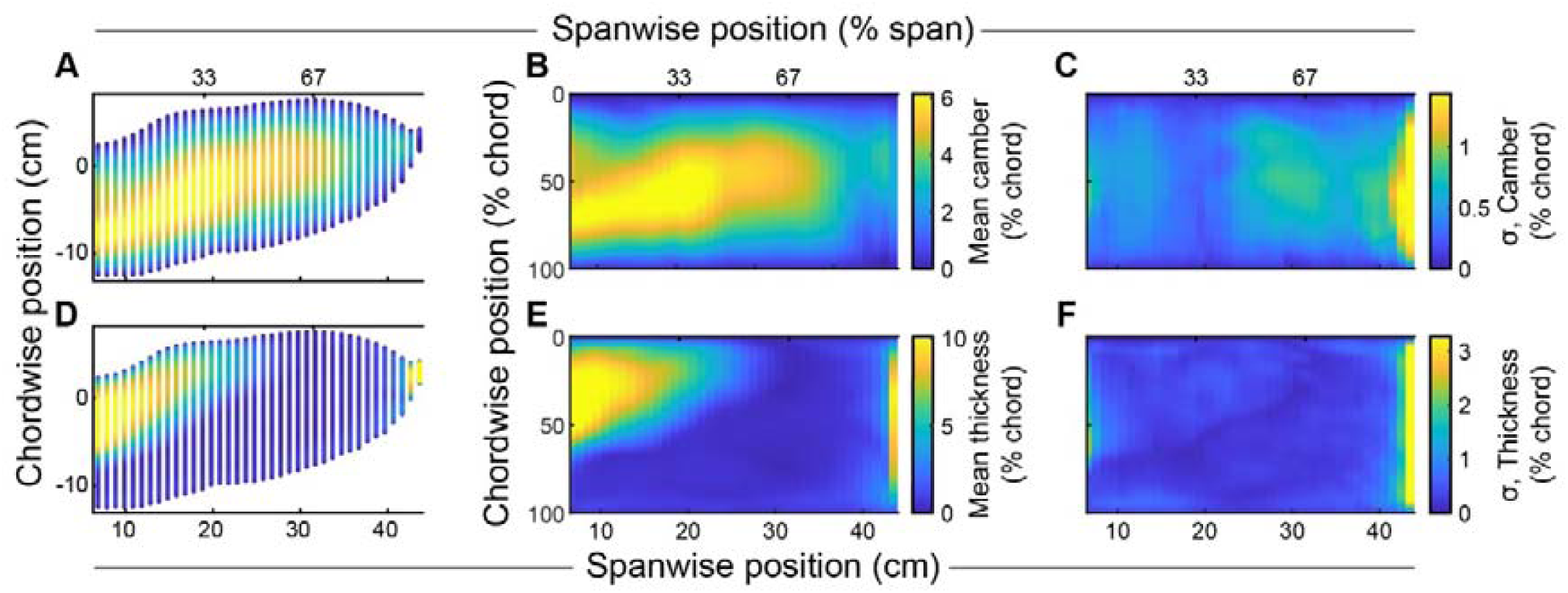
Wing planform distributions of camber (A-C) and thickness (D-F) are consistent across flights. A,D) Planform view of wing profiles, colour-coded by chord-normalized camber (A), and thickness (D). Normalizing chordwise position to percent chord (B,E) reveals that while thickness is generally anterior (E), peak camber (B) is posterior. (C,F) Standard deviation of camber (C) and thickness (F) across the planform are low.

**Figure 6.**
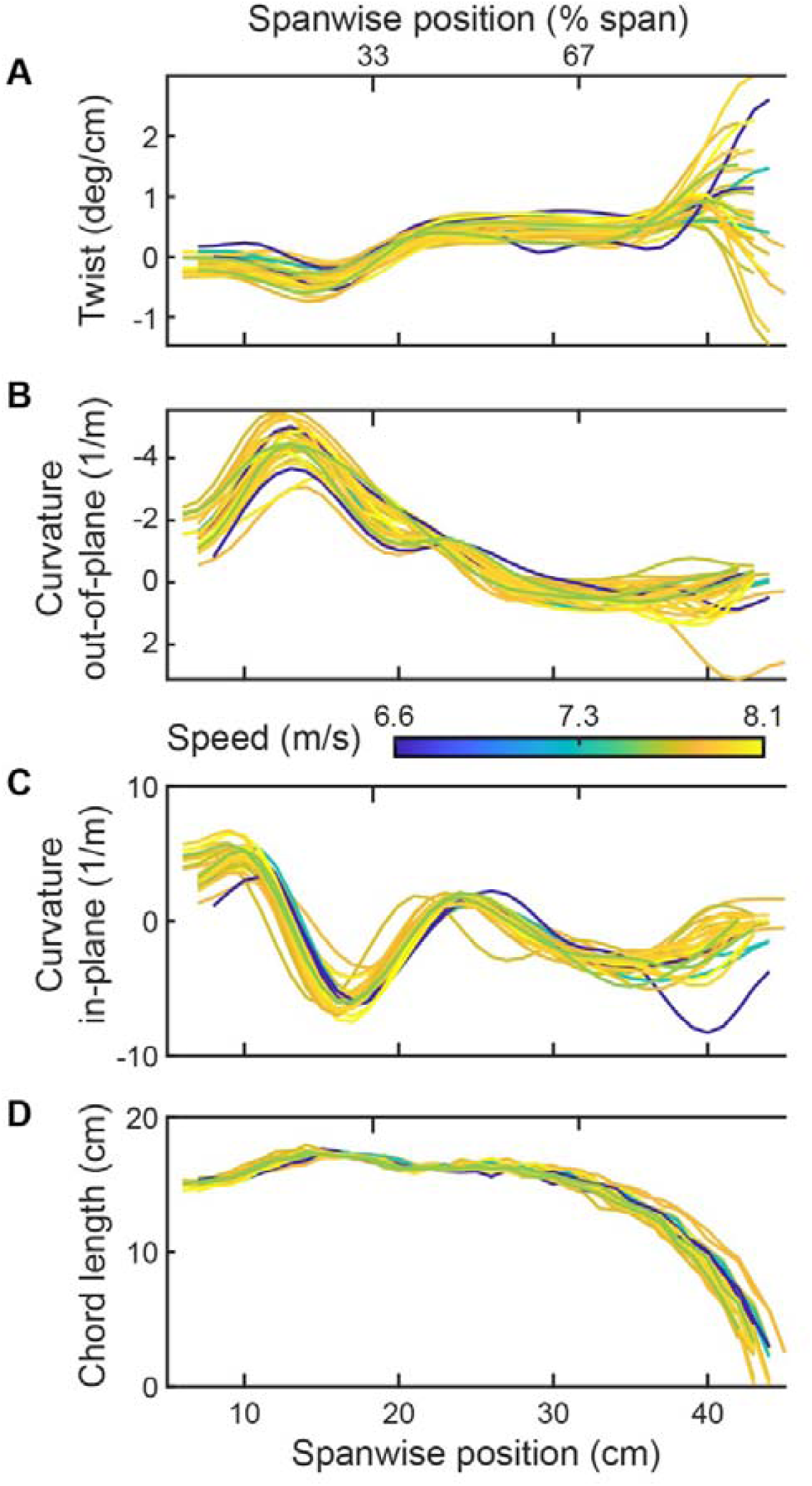
Wing section quantities are consistent across flights regardless of speed. Wing twist (A), curvature (B,C) and chord length (D) all follow the same pattern regardless of speed. Note, curvature out-of-plane (B) is inverted to provide a slightly more intuitive tracking of “spanwise camber”, as negative values are concave down.

### Wing shape

Wing camber is greatest in the proximal half of the wing, and peak camber gradually decreases in the distal half of the wing (Fig 5). Within the proximal half of the wing, peak camber along each chord is posterior or aft, occurring beyond 50% of the chord length. Maximum mean camber across the wing is 6.8% chord. Mean wing camber is positive throughout the wing.

The wing is thick in many regions, far more than just the regions that possess bones, specifically near the leading edge and proximally near the torso. Wing thickness exceeds 3% chord length for the majority of the chord when near the root (Fig 5). Distally, the wing is thin, as expected from anatomy. Near the root, we attribute the thickness throughout the chord to covert feathers. Near the leading edge, it is likely due to a muscularised, tissue flap-the propatagium. Maximum wing thickness is 11% of chord length.

Wing twist, which we describe as change in chord angle per unit span (given in degrees per cm), decreases and increases across the span (Fig 6A). From the root of the wing, near the shoulder--proceeding outboard--wing twist is negative and decreases in angle for the first third of the wing. After the first third of the wing length, a point coinciding roughly with the wrist, wing twist is generally positive. Wing twist is nearly constant from the wrist until the wingtip. This twist configuration is consistent with aerodynamic wash-in.

The quarter-chord position across the span is not a straight line, but a three-dimensional curve (Fig 6B,C, Fig 7). The quarter-chord curves both in and out of the plane of the wing, and we describe the movement as the curvature (1/r), as this makes our description less sensitive to the chosen coordinate system. Near the root, out-of-plane curvature is concave down (negative). At approximately mid-span, out-of-plane curvature becomes concave up, but remains near zero until the wingtip. In-plane curvature oscillates between forward sweeping and backward sweeping. Near the root, in-plane curvature sweeps the wing forward (positive) for the first 20% of the span. The next 20% span is strongly swept back, after which the curvature approaches zero until the wingtip.

**Figure 7.**
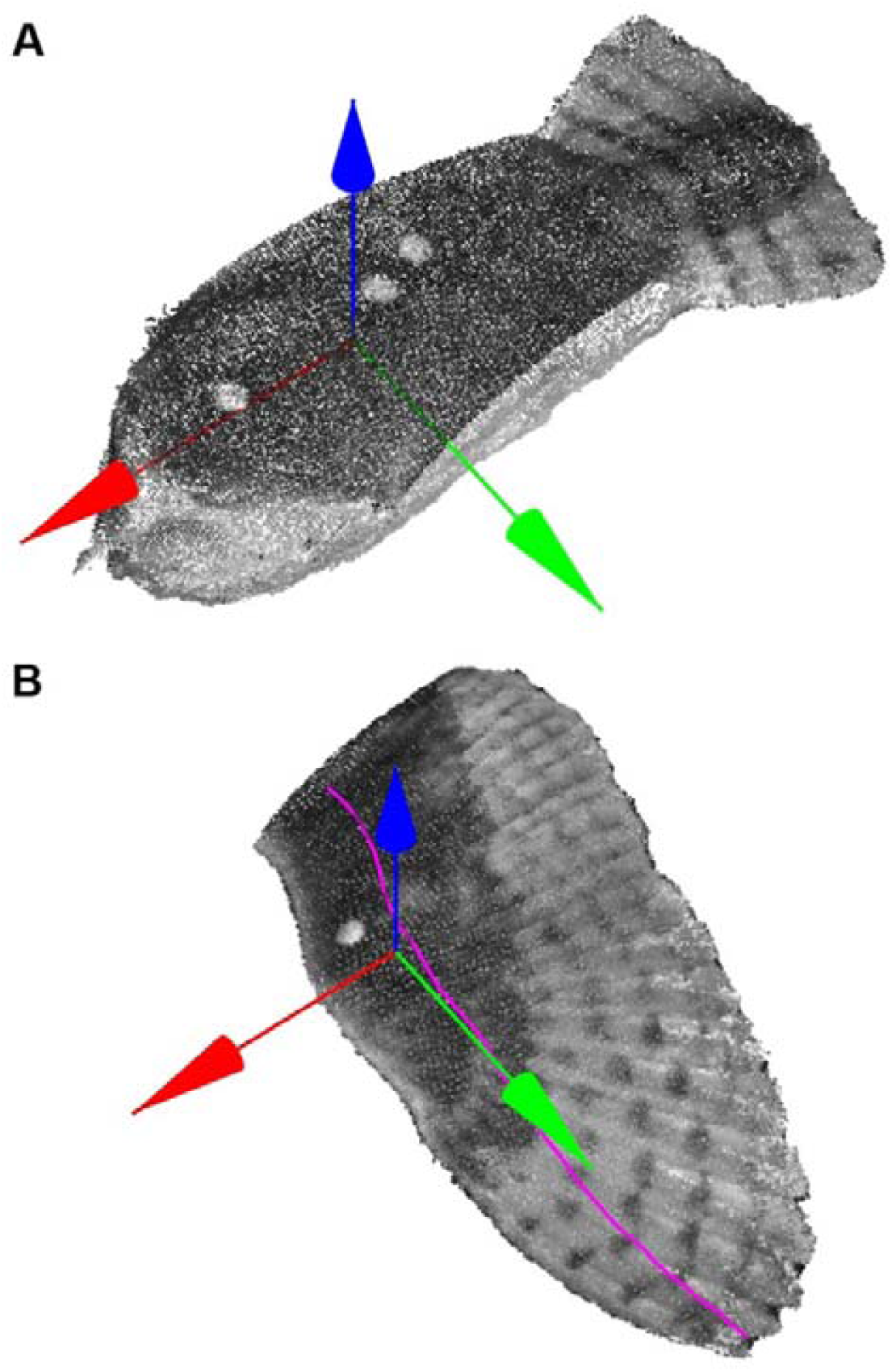
Body and wing coordinate system. (A) Elevated view of dorsal and left side of the head, torso, and tail. (B) Elevated view of the upper surface of the left wing. The purple line traces the quarter-chord position across the span. (A,B) Wing posture describes the rotations necessary to align the two coordinate systems.

### Relationship between speed and wing shape: camber, twist, and curvature

We tested for subtle relationships in wing shape but did not find any statistically significant or clearly aerodynamically significant effects. We tested for the effect of speed on three chord profiles: 25, 50, and 75% of maximum wing length. There was no relationship between speed and maximum camber, wing twist, or curvature in-plane or out-of-plane.

### Wing pose

While wing shape was essentially invariant among our flights, wing posture was not. Wing pitch—long-axis rotation of the wing—decreased with flight speed, but there was no relationship between sweep or roll and speed. Wing pitch decreased 0.7 ± 0.6 degrees for every meter per second increase in flight speed (r = -.24 ± .16; mean ± s.e.m). The relationships do not appreciably change if regressed against the square-root of flight speed, which is proportional to lift generated. Median wing pitch was 1.0 degrees up (range: -0.9 to 4.0).

Relative to the reference pose (Fig. 7), wing posture is swept back and depressed below the body in an anhedral configuration. Wings are swept backwards on average 3.7 ± 1.9 degrees (mean ± dev.). Wings form an anhedral angle on average of 7.0 ± 4.7 degrees (mean ± dev.).

### Testing our computational fluid dynamics model

We tested our computational fluid dynamic model by 1) comparing vertical force generation to weight support, and 2) by comparing the wake and downwash to those from the same individual gliding through tracked, neutrally-buoyant, soap bubbles. 1) After accounting for vertical acceleration, the vertical force of the CFD model accounted for 94.2% of weight support using the Shear Stress Transport (SST) model assumptions of flow. 2) The simulated downwash is in good agreement with the measured downwash both in magnitude and in distribution across the span (Figs. 8,9).

**Figure 8.**
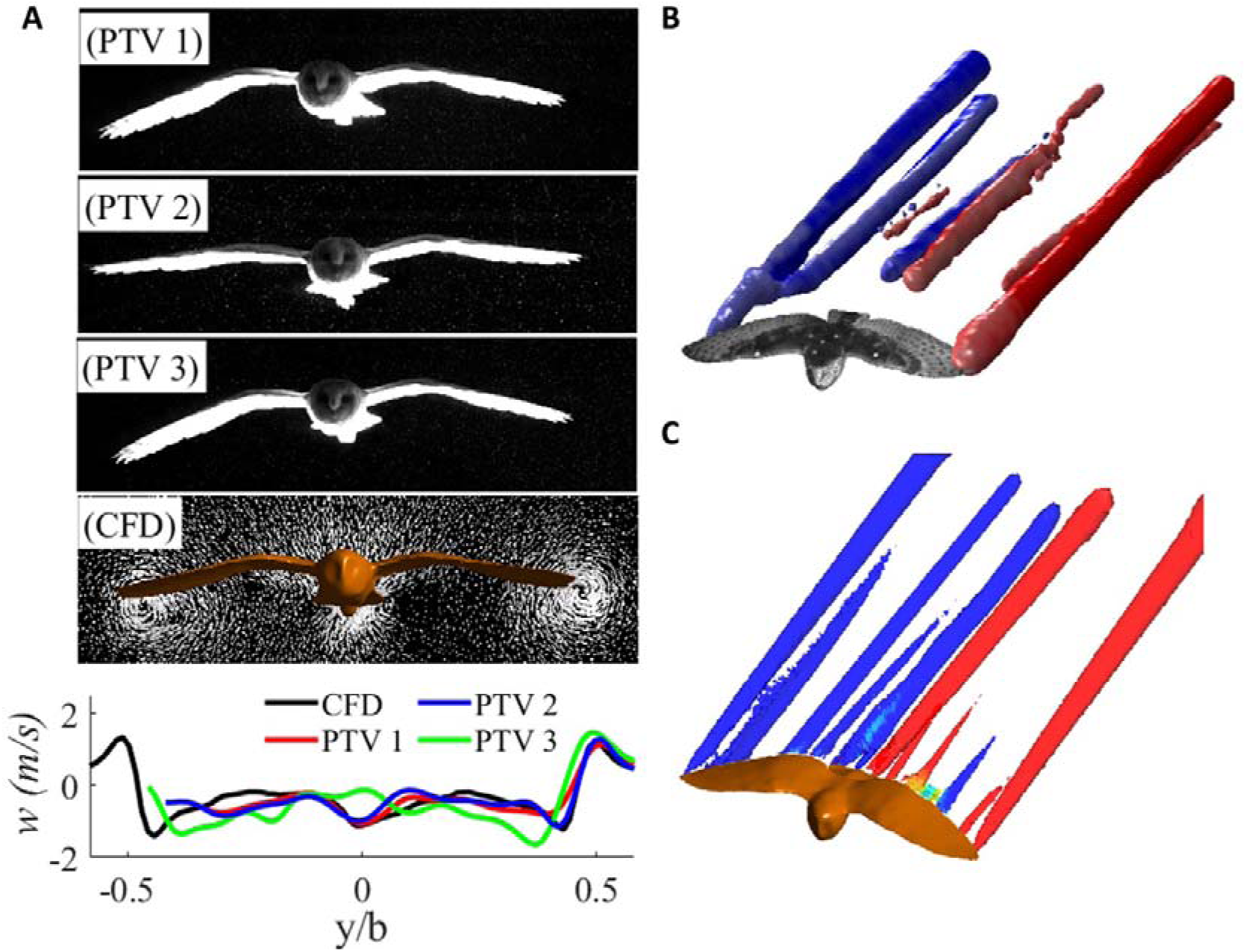
Downwash validation for a gliding T. alba. A) Projections of the posture adopted in each flight, along with the posture of the CFD surface model; and the corresponding measured and computed downwash in the transverse plane. B) The vortices in the wake measured using PTV (Q-value = 35 s^-2^; flight ‘PTV 1’) produce a complex wake pattern similar to that simulated (C). Note that the a) normalised downwash in CFD model (red) matches not only the measured magnitude across the wings, but also shares a similar distribution of downwash across the wing span. Not surprisingly, downwash behind the tail varies slightly.

**Figure 9.**
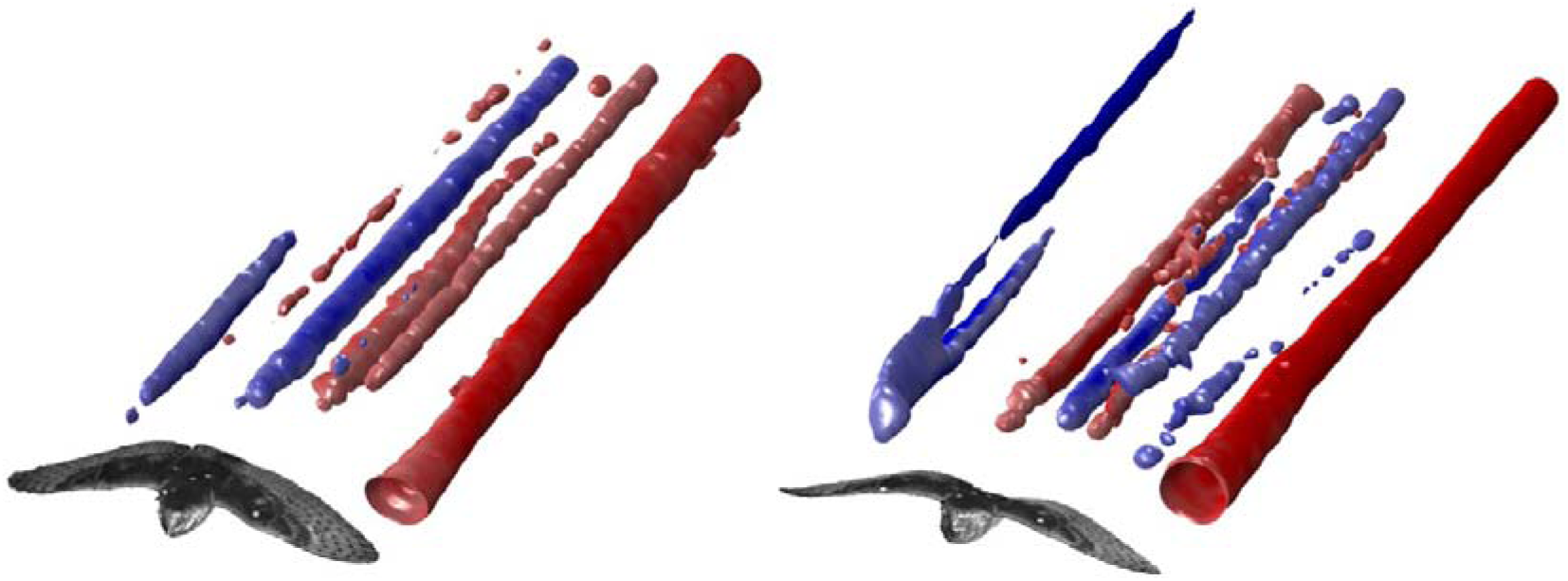
Additional views of the vortex structures measured using PTV (Q-value 35 s^-2^). Note, the right tip-vortex is not fully captured in the measurement. Left) ‘PTV 2’, Right) ‘PTV 3’

## Discussion

The results here 1) provides new insight into the relationship between morphing and flight speed; 2) details morphology of gliding wings at self-selected speeds; and 3) lays a foundation for future study of *T. alba* using computational fluid dynamics. While our range of flights speeds was limited to those self-selected, the wing shape used for the varying flight dynamics was consistent; and instead, wing pose, tail pose and tail shape varied with speed. This control strategy indicates that, at these speeds, *T. alba* is not using fine-wing shape control, but instead adjusts the wing’s pose and the tail’s shape and pose to modulate aerodynamic forces. Finally, we have identified a flight envelope for which we know 1) the 3D shape of the wing, 2) that this shape is repeatable, and 3) that the geometry can produce comparable spanwise distributions of downwash and vorticity to those measured using particle tracking velocimetry, when combined with an appropriate fluid mesh and turbulence model. We provide the geometrical 3D model of *T. alba*, to provide a strong foundation and to encourage others to explore gliding flight. Importantly, the wing shapes measured include aeroelastic effects and represent the actualised wing form, which likely differs from wing shapes measured without aerodynamic load. Note, that computational fluid dynamics using the model *can* replicate the correct fluid dynamics, but an appropriate fluid mesh is critical to proper computation.

### Relationships with speed

The patterns of wing and tail morphing among our flights are congruous with *T. alba* gliding relatively slowly. While published relationships between wing shape and speed lack similar detail, wingspan and area tend to change less at slow speeds (Tucker 1992), and birds consistently spread their tail in this speed range—consistent with what we observe (Tucker 1992; Rosen and Hedenström, 2001; Henningson and Hedenström, 2011). This slow-speed domain, as defined by the relationship between speed and wing and tail morphing, contains both the minimum-sink speed and maximum-range speed in *Corvus monedula*--jackdaw (Rosen and Hedenström, 2001).

We add to the published observations of wing shape and speed that, at slow speeds, wing shape is invariant. Wing camber, thickness, twist, quarter-chord curvature, and chord length follow a pattern across span and planform that does not vary with speed. Postural changes are the geometric angle of attack of the wing, and the pitch of the tail.

### Wing morphology

The general morphology of the wings is congruous with the patterns presented by Durston and colleagues (Durston et al., 2019). The specific individual studied, ‘Lily’, is the same in both studies. Our studies differ in that here we present indoor flight through quiescent air and perhaps because of this demonstrate a level of repeatability in wing shape previously unseen.

The measured wing morphology is not generalizable to other species. Wing twist is positive throughout the distal half of the wing and forms a wash-in configuration (Fig 10), as in *Aquila nipalensis* (steppe eagle; Carruthers et al., 2010), but not *Falco peregrinus* (peregrine falcon; Durston et al., 2019). A wash-out configuration is often used in conventional aircraft to avoid aerodynamic stall at the wingtips and increasing stability during a turn. Strong washout configurations can serve as yaw control and has been suggested as a means of manoeuvring in birds (Bowers et al., 2016).

**Figure 10.**
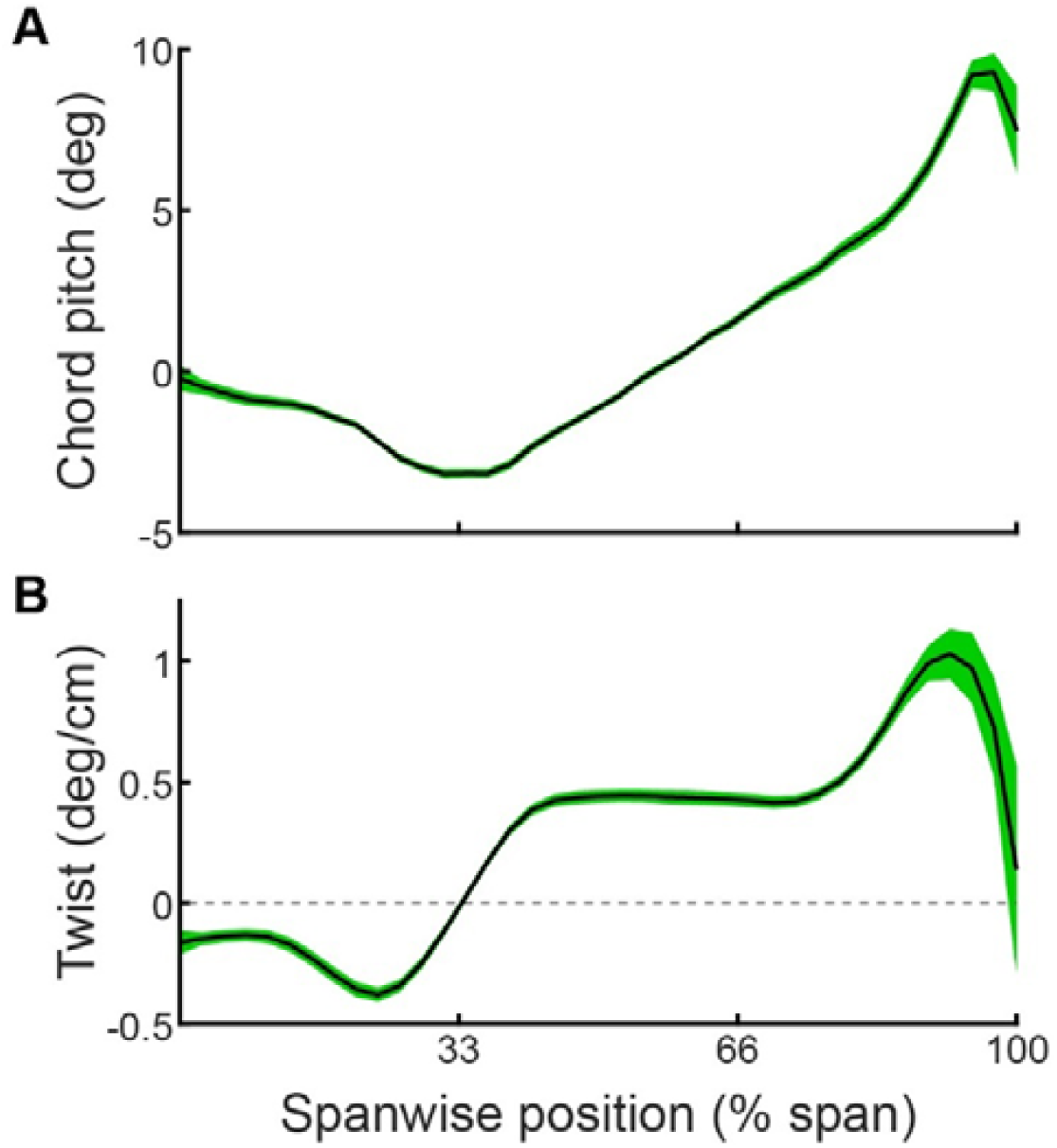
Spanwise distribution of pitch and twist among chords. Means (black lines) are bounded by standard error (green). Mean pitch (A) decreases slightly at the wingtips, but this is not observed when averaging twist (B). Mean twist remains positive across the distal region of the wing consistent with aerodynamic wash-in.

The camber profile of every chord section was positive across the chord (Fig 5). Aeroelastic interactions of feathers could have resulted in negative camber near the trailing edge (Carruthers et al., 2010), which can serve to enhance pitch stability by pulling the aerodynamic centre posterior/aft. Feathers are often treated as thin, cantilevered plates, and aerodynamic lift would produce a concave up deflection in a flat plate and local negative camber; however, feathers are curved along the rachis, and our results indicate for *T. alba* this curvature is sufficient-along with the feather’s bending stiffness-to resist any appreciable concave-up deflection.

### CFD and PTV

We found strong agreement between the downwash of three different flights and the computed downwash, despite the numerous potential sources of error from wake measurement, 3D reconstruction, 3D geometry generation, assumptions of similarity in wing pose and shape, and turbulence models which are not explicitly designed for this Reynolds number. From our kinematic results we believe that deviations between the measured and computed downwash are likely to be due to the assumptions of the CFD model and not variation in wing shape; noting however, we do not have simultaneous reconstructions of the 3D form of the bird and its wake. Where we observed variance in the measured wakes—and deviation from the CFD model—is the downwash behind the tail, which fits with our larger kinematic results, as tail-modulated lift is part of the control strategy for *T. alba* at these speeds (Fig 3).

### Individual versus species

An important caveat of this work is that our measured wing shape represents that of a single individual over a three-day period. Wing shape varies over life cycle, such as during moult (KleinHeerenbrink and Hedenström, 2016), and varies across individuals, although not enough data exist to determine the magnitude of variation in flight.

## Conclusions

*T. alba* does not use fine-wing shape control to glide at slow speeds in steady conditions, but instead relies upon wing postural control (gross pitch) and changes in both tail shape and pose to modulate aerodynamic force. Wing shape was invariant across all flights, despite dynamic pressure changing by 50% (speed by 23%). Wing shape invariance also makes having a three-dimensional model of the wing form a powerful tool for understanding the fluid dynamics of postural change and tail morphing in gliding flight. We provide this geometry, which we have tested against wake measurements and weight support, to assist other researchers interested in exploring the fluid dynamics behind gliding flight. It is interesting that despite the dramatic morphing that occurs during flapping flight, that *T. alba* utilizes a constant wing shape over our fifteen different glides and we expect future work to address whether this is primarily a constraint on morphing/control or one of fluid dynamic optimisation.

## Methods

### Bird

The study subject was a captive-bred, adult female *Tyto alba* (barn owl). The bird was trained to fly between handlers on command and land on a perch where she received a food reward after each flight. All work was approved by the Ethics & Welfare Committee of the Royal Veterinary College (URN 2015 1358) and the University of Bristol Animal Welfare and Ethical Review Body (UIN UB/15/070) and complied with all relevant ethical regulations.

### Experimental flights

Flight measurements were conducted indoors in a 17 m long flight corridor at the Royal Veterinary College (Hatfield, UK). The corridor was 2 m wide, bounded by a structural wall on one side and framed mesh on the other, with a suspended floor to elevate the flight path of the bird and permit camera views from above and below. Each flight consisted of early flapping to build speed, prior to entering a smooth glide when the measurement was taken, and finally perching 3-5 m beyond the point of measurement.

### Wing surface measurement

We imaged the bird using an array of twelve, synchronized, high-speed cameras, arranged in upper and lower sets (Fig 1), comprising pairs of either Photron FASTCAM SA3 (1024×1024 pixels), FASTCAM SA-Z (1024×1024 pixels) or Mini WX100 (2048×2048 pixels) models (Photron Europe Limited, West Wycombe, UK). Camera placement ensured that all cameras viewed either the ventral or dorsal surface when the bird was in the centre of the measurement. A cross-section through the measurement volume at gliding height was approximately 2 × 2 m.

Cameras recorded at 500 frames per second and illumination was provided by custom stroboscopic LED lamps. The combination of short exposure periods of ∼1/3000 s and moderate frame rates allowed the LEDs to operate at a ∼17% duty cycle, reducing average illumination intensity and heat production. Where possible the imaging background was covered in black material to facilitate automated masking of the bird during later image processing.

Birds were marked with motion-capture retroreflective markers. Motion capture was not used in the analysis presented here. The markers are visible in the reconstructions as white discs.

Three-dimensional surface points of the bird were reconstructed using commercial photogrammetry software (Photoscan version 1.3.5; Agisoft LLC, St Petersburg, Russia) and custom Python scripting. Within the Photoscan pipeline, first, common image features are identified and matched between multiple views, providing an initial sparse reconstruction, when combined with camera calibrations. The sparse reconstruction then serves as a foundation for disparity map calculations between camera pairs. The 3D point cloud is then reconstructed from disparity maps and camera calibrations. Each cloud point is assigned a grayscale value, based on the matched image pixels from which it is obtained.

### Camera Calibration

Camera calibration involved three steps: (i) intrinsic calibration; (ii) individual extrinsic calibration of upper and lower cameras sets; and (iii) alignment of both sets’ coordinate systems to the corridor reference frame. (i) The intrinsic properties of each camera-lens pair, including optical distortion, were calculated from 50 to 100 images of a flat, 1.2 × 0.7 m checkerboard that filled the field of view of each camera. To fill the view and achieve a sharp image, we increased sharpness over a larger depth of field by further closing the aperture; we assumed changing aperture had negligible effects on the intrinsic calibration. We did not change focus, as that would change the focal length of the lens. (ii) The extrinsic parameters of each camera set--camera positions, orientations, and scale--were calculated from images of a visually-textured board with corner markers to define scale. (iii) As the pattern on the board could only be seen from a single set of cameras at a time, we aligned the upper and lower sets of cameras using images of a T-shaped wand with spherical reference points. Finally, the coordinate system was set using an L-shaped wand, directed along the flight corridor and levelled in the plane normal to gravity.

### Estimating camera calibration accuracy

Reconstruction error can be attributed to two sources, error in camera calibration and error in point matching. To estimate camera calibration error, we compared point cloud surfaces generated of a physical model to a high-accuracy laser-scan (Romer Absolute Arm, RA-7525-SI, accuracy 0.063 mm). To generate point cloud surfaces, we reconstructed a visually-textured fibreglass model of a bird placed within the measurement volume at glide height. Visual texture was added to the model using a marker to make small dots of varying size, which enhances effective point matching. Model wingspan was similar to the barn owl. Error in point matching depends upon the sharpness and size of the matched features on the surface, which for the bird is feather colour and texture, which varies across it.

### Criteria for exclusion

We excluded any flight where the wing was moving relative to the body, as our analysis of a single instance in flight would be an inaccurate portrayal of wing shape and pose. This removed two flights of our seventeen.

### Geometry processing – cleaning

In general, point clouds computed through disparity maps contain noise due to spurious false-positive matches. We minimised false positives through image masking by performing background subtraction against the temporal-mean background image. We removed remaining false positives manually.

### Wing and body posture - markerless alignment

To calculate changes in posture, *i.e*., rigid-body transformations, we did not use markers, but instead aligned the complete three-dimensional shapes (tens of thousands of points) of the segmented wings or body to a reference shape. Alignment was performed using an iterative closest point alignment algorithm (as implemented in Matlab 2019a: pcregistercicp.m). This reference shape was then oriented to a reference-coordinate system to make the postural transformations intuitive.

The reference body and wing shapes came from a single flight--we used the left wing as the reference shape for both wings, due to its slightly higher point density. Our selected flight was based upon its dynamics: low overall acceleration, average body pitch, low body-yaw relative to the self-induced flow, and average speed. The similarity in wing shapes and body shapes (not shown) across flights meant that the exact chosen flight was relatively unimportant.

The body reference pose was defined by two vectors. A vector running from the base of the tail to the tip of the beak formed the anterior-posterior (fore/aft) axis. The lateral axis used to define body pitch was orthogonal to the plane defined by the anterior-posterior and gravitational axes. Finally, the axis defining body yaw was orthogonal to the lateral and anterior-posterior axis. We discuss our sign conventions for rotations below.

We did not compute the roll component of body pose across flights. This was due to the difficulty in resolving rotations about the anterior-posterior axis. The body is nominally a tube, and the most dramatic alignment features were the feet and the wing roots, which would not obey the assumption of rigid-body movement.

The wing reference pose was nominally flat. We fit a plane to the left wing to correct for long-axis rotation/pitch; as well as wing elevation/’dihedral’. Once these rotations were corrected for, this placed the wing in an arbitrary, but biologically and aerodynamically relevant, swept position (Fig 7). This swept posture is 3.7 degrees swept forward from the average of all wings, and if tracking quarter-chord position, 1.8 degrees swept forward at the root, and 5.6 degrees swept forward from wing-tip to wing root.

### Rotational movements

We deconstructed our rotation matrices into Euler angles using different, but fundamentally similar, sets of rotations for the wing and body. We calculated the rotations about the body in the order of yaw, pitch, and then roll. Long-axis rotation of the body— roll—is similar to long-axis rotation of the wing—pitch; so we adjusted the rotation order for the wings to be sweep, elevation/’dihedral’, and then pitch. The axis order would have been analogous if the long-axes for the reference postures of the wing and body were parallel.

We defined positive upward pitch for the body, wings, and tail as upward movement of the anterior edge and downward movement of the posterior edge. Sweeping the wing backward, by placing the wingtip further posterior, and elevating the wing, moving the wingtip away from the ground, were both positive rotations. We defined glide angle to be positive if the bird’s downward velocity was negative.

### Geometry processing – wing and tail segmentation

By aligning the torso to the same coordinate system, we could segment the wings, tail, and head, using consistent spatial criteria. We used a custom-written interface to manually define the spatial volumes segmenting the bird.

### Geometry processing – wing chord profiles

Wing chord profiles were defined as cross-sections in the wing-reference pose (Fig 7) orthogonal to the lateral axis (wing pitch vector). Spacing between chords was 1 cm along the span. We defined the trailing edge point as the posterior/aft most point within 0.1 cm of the defined spanwise location of the profile. The location of the leading edge was the point with greatest curvature within the anterior region of the wing. To compute curvature, we fit a polynomial surface to the points within 4 cm spanwise of the profile. The polynomial surface was semi-parametric and used a moving window to determine which points were fit. The window moved along the lower surface of the wing, wrapped around the leading edge, and then continued along the upper surface to the trailing edge. The window size contained points within 4 cm spanwise of the profile, and 4% of the points anterior and posterior of the chordwise point on the surface. The polynomial surface was a linear fit along the spanwise axis, and a fourth-order polynomial along the chord axis. This computational approach also provided some smoothing of the wing surface.

We computed the mean line using the proportional distance travelled along the lower and upper surface from leading to trailing edge. After computing the upper and lower surface length, we discretised the surface into evenly spaced percentage of distance travelled. The mean of the two surfaces at each discretised percentage determined the mean line.

The quantified wing shape parameters were assessed using the leading and trailing edge points, the mean line, the quarter-chord position of each chord profile, and the two surface fits. Chord pitch described the angle from leading edge to trailing edge within the plane of the profile. Camber described the pitch-corrected displacement of the mean line across the profile. Quarter-chord was determined as one-fourth the linear distance from the leading edge to the trailing edge. Wing thickness described the distance between the surfaces at proportional distances along each surface.

### Geometrical model of the bird

The geometrical model was formed manually using PhotoScan to develop the mesh of each surface, meshlab to combine the surfaces and smooth, and finally conditioning the geometrical model to be appropriate for CFD analysis using Materialise Mimics (Materialise NV, Leuven, Belgium) and Ansys SpaceClaim (Version 19.1, ANSYS, Inc., Canonsburg, PA, USA).

### Computational Fluid Dynamics

The fluid mesh, built around the bird surface, was generated in ANSYS Meshing (Ansys Inc., Canonsburg, USA). The simulation domain (domain size: 9000 × 6000 × 6000 mm) was discretized by approximately 15 million non-uniform volume elements. Two bodies of influence controlled mesh size: mesh size was 5 mm for the inner body of influence and 12 mm for the outer body of influence. Within the inner body, adjacent to the bird surface, 15 inflation layers were used to resolve the boundary layer: the first layer thickness was 0.5 mm (y^+^ ∼10) and subsequent layers had a growth ratio of 1.2. We found that if we further refined the near-surface mesh by decreasing the first layer thickness to 0.1 mm (y^+^ ∼3), there was only a 3% difference in the lift of *T. alba*.

Flow around the bird surface was computed using the k-ω SST turbulence model in ANSYS Fluent (version 19.1; ANSYS Inc., Canonsburg, USA). The SST model uses a blending function which combines the merits of the standard k-ω model, for near-surface simulation, and the standard k-ε model, for the domain away from the surface, which is appropriate for bird flight simulations that require a high accuracy boundary layer.

#### CFD validation using PTV

To validate our CFD simulations, we used particle tracking velocimetry (PTV; Lavision UK, UK) to measure the wake of a gliding barn owl. The owl flew between handlers through a corridor that provided a dark background, enhancing the contrast of helium-filled soap bubbles illuminated by an array of white LEDs. The corridor was approximately 2 m wide, 1.8 m high and 14 m long. The measurement volume, located at the exit of the corridor, was illuminated only after the bird had entered it. We filmed with eight, high-speed, synchronised cameras but found the optimal number to be four, with which we were able to track on the order of 20,000 neutrally-buoyant, soap bubbles of 0.3 mm diameter. Further details of the PTV experiment are described in Usherwood *et al*. (2020).

## Supporting information

Tyto alba geometry

## Acknowledgements

We thank Nathan Phillips for helpful discussions and assistance during setup and data collection, and Maja Lorenc for assistance during data collection. We also thank Lloyd and Rose Buck for their falconry expertise during the flight testing. This project has received funding from the European Research Council (ERC) under the European Union’s Horizon 2020 research and innovation programme (grant agreement No 679355). This material is based upon work supported by the Air Force Office of Scientific Research, Air Force Material Command, USAF under Award No. FA9550-16-1-0034. JRU funded by a Wellcome Trust Fellowship 202854/Z/16/Z.

## Notes

### Competing Interest Statement

The authors have declared no competing interest.

